# A look into the future: Using a transcriptomic meta-analysis of *Diptera*-*Wolbachia* systems to project the sustainability of arboviral control strategies

**DOI:** 10.1101/2023.09.03.556149

**Authors:** Sebastián Mejías, Natalia E. Jiménez, Carlos Conca, J. Cristian Salgado, Ziomara P. Gerdtzen

## Abstract

**Background:** An effective strategy for arboviral control consists in transfecting *Aedes aegypti* mosquitoes with the intracellular bacteria *Wolbachia pipientis*, which reduces host viral susceptibility and spreads itself into wild populations via reproductive manipulations. However, the prospect of losing the efficacy of this strategy underscores the need for deepening the mechanistic knowledge of *Diptera*-*Wolbachia* systems and identifying relevant *Wolbachia* effects that could decline upon adaptation of *A. aegypti* transfections. A systematic comparison of publicly available *Diptera*-*Wolbachia* transcriptomic datasets could yield progress in this matter.

**Methodology/Principal findings:** We derived differentially expressed gene (DEG) sets from previously published *Diptera*-*Wolbachia* transcriptomic datasets, subjected them to enrichment analysis of Gene Ontology terms, and intersected the results to identify patterns of host gene/function regulation by *Wolbachia*. A putative farnesoic acid methyl transferase (*AAEL004667*) and a flavin-containing monooxygenase (*AAEL000834*) were consistently upregulated in transfected *A. aegypti* and linked to cytoplasmic incompatibility and viral susceptibility, being proposed as novel targets of study. Genes implicated in viral blocking *—GNBPA1*, *PGRPS1*, *DEFC*, *Tf1*, serine-type endopeptidases and endopeptidase inhibitors— were consistently upregulated in transfected *A. aegypti* but not in native infections, indicating that they could lose responsiveness to *Wolbachia* over time and should be considered to keep the efficacy of arboviral control. The commonality of chitinase regulation by *Wolbachia* was identified and proposed as an explanation for the loss of desiccation resistance in transfected *A. aegypti*’s eggs, which is a main obstacle for the introgression of *Wolbachia* in mosquito populations.

**Conclusions/Significance:** The present work points out relevant gene targets to consider for arboviral control sustainability and provides new hypotheses for deepening the understanding of *Diptera*-*Wolbachia* systems.

**Author Summary:** Arboviral diseases (e.g. dengue), which are mainly transmitted by the mosquito *Aedes aegypti*, impose a global public health crisis. An effective strategy for controlling the spread of these diseases is to artificially infect *A. aegypti* populations with the bacteria *Wolbachia pipientis*, which reduces its capacity to transmit arboviruses. However, future adaptive changes in the novel *A. aegypti*-*Wolbachia* association could diminish the efficacy of this approach. To prevent this, it is crucial to have a solid biological understanding of *Wolbachia* infections and predictions about specific changes that artificial infections could undergo. By analyzing publicly available biological data from *Wolbachia*-infected mosquitoes and flies we were able to propose new hypotheses regarding general aspects of *Wolbachia* infection and to identify antiviral effects of *Wolbachia* in *A. aegypti* that could decline over time, thus providing relevant information for keeping sustainability of a key arboviral control strategy.

## Introduction

Arboviruses (<u>ar</u>thropod-<u>bo</u>rne <u>viruses</u>) such as Dengue, Zika and Chikungunya (DENV, ZIKV and CHIKV, respectively) impose serious morbidity and mortality burdens worldwide, which have been on the rise over the last two decades [1–8]. Arboviral vectors, mainly *Aedes aegypti* and to a lesser extent *Aedes albopictus*, have spread and are now present in Africa, Europe, Asia, Oceania and the Americas [3,5,9]. This together with the general lack of effective broad-spectrum vaccines and the weaknesses of insecticide-based vector control have driven the development of new strategies against arboviral diseases [2,7,10–12].

One such novel approach is based on artificially infecting (*transfecting*) arboviral vectors with *Wolbachia pipientis*, an intracellular, maternally inherited bacteria naturally found as an endosymbiont in nearly half of insect species [2,13]. *Aedes aegypti* mosquitoes can be transfected to yield individuals with arboviral resistance (pathogen or viral blocking) and reproductive advantages with respect to their wildtype [14–16]. These traits have been exploited by releasing transfected mosquitoes that spread *Wolbachia* and lower the risk of disease for humans due to the viral blocking phenotype. Indeed, stable *Wolbachia* introgression in wild mosquito populations and subsequent arboviral disease burden reduction has been achieved in Indonesia [17], Australia [18] and Brazil [19], posing it as a promising control approach.

Despite its current success, there are concerns about the long-term efficacy of this strategy, particularly regarding future phenotypic changes of transfected mosquitoes [10,11,13,20,21]. These issues are sustained by observation of native *Wolbachia* infections, which are deemed as reflective of the future state of transfections once a host-*Wolbachia* coadaptation has taken place in wild conditions [13,20]. Native infections tend to show considerably lower viral blocking, bacterial density, virulence and tropism [13,22–25]. In fact, substantial phenotypic changes of *Diptera*-*Wolbachia* systems in the wild can occur in the scale of years [26]; even a complete shift from deficit to enhancement of fecundity by *Wolbachia* has taken place in just two decades [27]. Hence, acquiring mechanistic knowledge of both native and artificial host-*Wolbachia* systems is crucial to preserve the efficacy and safety of this strategy, to prevent or react to possible future events such as attenuation of viral blocking [13,21,28].

Multiple transcriptomic studies have compared gene expression in *Wolbachia* infected and uninfected hosts, showing effects of *Wolbachia* on diverse processes such as reproduction [29,30], redox homeostasis [13,26,31–34], sensory perception [13,26,29], metabolism [13,26,28,29,31,32], proteolysis [26,31,32,34] and innate immunity (e.g. the Toll pathway) [26,29–32,34,35]. The wide variety of the reported effects opens an essential question: are there transcriptomic signatures of *Wolbachia* across biological contexts? To the best of our knowledge, the only study dedicated to such pattern search was the transcriptomic meta-analysis performed by Chung et al. (2020) [36], although strictly focusing on the bacterial transcriptome and mainly on filarial nematodes.

In this study we performed a meta-analysis of *Diptera*-*Wolbachia* transcriptomic datasets published in the last two decades to identify patterns in *Wolbachia*’s transcriptomic effects across different contexts. In particular, to broaden general mechanistic knowledge of *Wolbachia* we searched for transcriptomic effects that are common across *Diptera* systems that would likely reflect fundamental aspects of their endosymbiosis. To predict future changes on *A. aegypti*-*Wolbachia* that are relevant for the sustainability of arboviral control, we focused on effects that differentiate this transfected system from native infections, as these last ones serve as a proxy for the coadapted transfected state.

## Methods

### Overview

Transcriptomic datasets comparing *Wolbachia*-infected/uninfected *Diptera* conditions were retrieved from public databases and analyzed to obtain differentially expressed gene (DEG) sets. Previously published DEG sets were used directly only when raw data was unavailable. To develop a generalized transcriptomic profile of *Aedes aegypti*-*Wolbachia*, DEGs that appeared in at least 7 of 8 DEG sets from this system were selected.

Furtherly, to find patterns of *Wolbachia* effects on gene expression across *Diptera* hosts, all DEG sets were mapped to *Drosophila melanogaster* orthologs and intersected. To find common effects of *Wolbachia* on *Diptera* host’s functions, all DEG sets were subjected to enrichment analysis of Gene Ontology (GO) terms and the five most frequently enriched terms were selected. Methodological details are provided in the following.

### Building DEG sets from microarray data analysis

Raw microarray two-channel intensity data was retrieved from Gene Expression Omnibus [37] or ArrayExpress [38] databases. All analyses were performed using R v4.2.3 [39] with the limma v3.54.2 package [40], and default parameters and standard methods were used unless stated otherwise. To prevent unreliable microarray spots from affecting normalization steps, weights of outlier and quality flagged spots were set to zero. After background correction, two-channel data was combined into log-ratio values and within-array normalization was performed applying the global *loess* method [41] in case of non print-tip based arrays. Then, between-array normalization was performed and control probes were removed before differential expression analysis, considering average intensities for duplicated spots. DEGs were defined as those with expression changes significantly greater than 20% (i.e., those rejecting H_0_: |β| < log(1.2) with adjusted *p*-value < 0.05, where β is the log-ratio of infected/uninfected mean expressions), a threshold that lies within a reasonable range for finding biologically meaningful effects [42].

### Building DEG sets from RNA-Seq data analysis

Raw RNA-Seq reads were retrieved from Sequence Read Archive [43] or Gene Expression Omnibus [37] databases. All analyses were based on a reference genome except for *Aedes fluviatilis* data, which required a *de novo* transcriptome assembly. RNA-Seq analysis tools were accessed from Galaxy v.23.0 [44] and used with default parameters, unless otherwise stated.

### Analysis based on a reference genome

Quality of raw *Drosophila melanogaster* reads was assessed with FastQC v0.11.9 [45] and then Trimmomatic v0.38 [46] was used to trim adapter and low quality sequences, using a sliding window approach. STAR v2.7.8a [47] was used to align trimmed reads to the *dm6* reference genome of *D. melanogaster* [48], with the splicing event information provided by the FlyBase annotation *dmel_r6.32* [49]. Aligned reads were counted at the gene level using featureCounts v2.0.1 [50] and differential expression analyses were done with DESeq2 v1.34.0 [51] implementation in R. A differential expression threshold of 20% expression change (adjusted *p*-value < 0.05) was established to identify DEGs, as in the microarray data case.

### Analysis based on a de novo transcriptome assembly

Raw *Aedes fluviatilis* reads quality was assessed with FastQC. Obtained reads were then subjected to two separate trimming procedures: one for transcriptome assembly and one for alignment to the assembled transcriptome, this last one being as described previously (Section 2.3.1). For transcriptome assembly, raw reads were subjected to additional Trimmomatic operations to trim low-quality outer bases and to discard short reads or overall low quality reads, so as to improve assembly quality [52]. After verifying trimming results with FastQC, the *A. fluviatilis* transcriptome was assembled using Trinity v2.9.1 [53], and its completeness was assessed by searching for *Diptera* benchmarking universal single copy orthologs with BUSCO v5.3.2 [54]. RSEM v1.3.2 [55] was used for read alignment to the assembled transcriptome and to estimate counts at the gene level, being provided with the gene-to-transcript map generated by Trinity. Estimated gene counts were then subjected to normalization and differential expression analysis with DESeq2, as described previously (Section 2.3.1). Obtained DEGs were annotated by searching for open reading frames (ORFs) with Transdecoder v5.5.0 [56], representing each DEG by its most highly expressed transcript (i.e., the one with greater mean count across all samples). Given that *A. aegypti* annotation is more curated than the one from *A. fluviatilis* [13], predicted ORFs were subjected to BLASTp [57] searches with default parameters against *A. aegypti* protein sequences kept in the VectorBase database release 62 [58], and DEGs inherited the annotation of their corresponding best BLAST hit.

### Functional enrichment analysis

g:GOSt module of g:Profiler version e109_eg56_p17_1d3191d [59] was used to perform enrichment analyses of Gene Ontology (GO) functional terms, using Fisher’s one sided exact test. Up and downregulated gene sets were analyzed separately, as this has been shown to improve statistical power for finding biologically relevant functional effects [60] and allows to distinguish preferent directions of regulation. For testing any given RNA-Seq-derived DEG set, the background gene set was defined as all annotated genes in the corresponding reference organism, while for each microarray-derived DEG set it was defined as all annotated genes that were represented in the corresponding array. *p*-values were adjusted by the Benjamini-Hochberg method to account for multiple hypothesis testing.

### Orthologous gene mapping

In order to allow a direct comparison of *Wolbachia* effects at the gene level for different host species, all DEG sets were mapped to *D. melanogaster* orthologs using the g:Orth module from g:Profiler. For the analysis and discussion of results, obtained g:Profiler orthologous assignments were compared with those of the OrthoDB database [61].

## Results and discussion

### Obtained DEG sets

A total of 18 DEG sets were obtained from 4 microarray and 7 RNA-Seq datasets published between 2009 and 2023. The collected data comprised transfected systems (*A. aegypti*-*w*Mel and -wMelPop) and naturally infected systems (*A. fluviatilis*-*w*Flu, *D. melanogaster*-*w*Mel and *D. paulistorum*-*w*Pau), as well as multiple tissues (heads, abdomens, testes, ovaries, muscles and whole bodies). Both female and male host data are considered, although the latter was represented in only 4 DEG sets, corresponding to natural infections. All samples correspond to adults, with ages varying from 1 to 15 days post eclosion [13,28,29,31–35,62], with the exception of larval testes sampled by Zheng et al. (2011) [36]. Table 1 summarizes the origin of all DEG sets together with the identifiers that will be used to refer to them here on.

**Table 1.** Summary of analyzed DEG sets.

| Set ID | Infection type | Host species | Strain | Tissue | Sex | Datatype | Accession number | Article (date) |
| --- | --- | --- | --- | --- | --- | --- | --- | --- |
| Aa1 | Transfection | *A. aegypti* | wMel | Whole | f | Array | E-MEXP-2931 | Rancès et al. (2012) |
| Aa2 | Transfection | *A. aegypti* | wMelPop | Whole | f | Array | E-MEXP-2931 | Rancès et al. (2012) |
| Aa3 | Transfection | *A. aegypti* | wMelPop | Whole | f | Array | PRJNA118709 | Kambris et al. (2009) |
| Aa4 | Transfection | *A. aegypti* | wMelPop | Muscle | f | Array | E-MEXP-2907 | Ye et al. (2013) |
| Aa5 | Transfection | *A. aegypti* | wMelPop | Head | f | Array | E-MEXP-2907 | Ye et al. (2013) |
| Aa6 | Transfection | *A. aegypti* | wMel | Whole | f | RNASeq | PRJNA867516 | Wimalasiri-Yapa et al. (2023) |
| Aa7 | Transfection | *A. aegypti* | wMel | Whole | f | RNASeq | PRJNA867516 | Wimalasiri-Yapa et al. (2023) |
| Aa8 | Transfection | *A. aegypti* | wMel | Whole | f | RNASeq | PRJNA867516 | Wimalasiri-Yapa et al. (2023) |
| Af | Native | *A. fluviatilis* | wFlu | Whole | f | RNASeq | PRJNA320882 | Caragata et al. (2017) |
| Dm1 | Native | *D. melanogaster* | wMel | Testes | m | Array | PRJNA147175 | Zheng et al. (2012) |
| Dm2 | Native | *D. melanogaster* | wMel | Testes | m | RNASeq | PRJNA639180 | Dou et al. (2021) |
| Dm3 | Native | *D. melanogaster* | wMel | Whole | f | RNASeq | PRJNA682591 | Lindsey et al. (2021) |
| Dm4 | Native | *D. melanogaster* | wMel | Whole | f | RNASeq | PRJNA602188 | Detcharoen et al. (2021) |
| Dm5 | Native | *D. melanogaster* | wMel | Ovaries | f | RNASeq | PRJNA439370 | He et al. (2019) |
| Dp1 | Native | *D. paulistorum* | wPau | Head | m | RNASeq | PRJNA515416 | Baião et al. (2019) |
| Dp2 | Native | *D. paulistorum* | wPau | Abdomen | m | RNASeq | PRJNA515416 | Baião et al. (2019) |
| Dp3 | Native | *D. paulistorum* | wPau | Head | f | RNASeq | PRJNA515416 | Baião et al. (2019) |
| Dp4 | Native | *D. paulistorum* | wPau | Abdomen | f | RNASeq | PRJNA515416 | Baião et al. (2019) |

E-MEXP prefix corresponds to ArrayExpress accession numbers, while PRJNA prefix corresponds to SRA and GEO accession numbers. DEG set identifiers were assigned according to host species. Aa: *Aedes aegypti*. Af: *Aedes fluviatilis*, Dm: *Drosophila melanogaster*, Dp: *Drosophila paulistorum*, f: female, m: male.

All datasets except for those from Baião et al. (2019) [29] and Wimalasiri-Yapa et al. (2023) [26] were analyzed *de novo*, yielding reliable results according to intermediate step quality assessments. Remarkably, for all RNA-Seq samples the trimming and mapping steps proceeded with >96.4% and >94.8% retained and mapped reads (or read pairs), respectively. *A. fluviatilis* assembled transcriptome represented 90.3% of *Diptera*’s benchmarking universal single-copy orthologs (85.3% in a complete form), comprising 63,416 contigs with median size of 550 bp and N50 equal to 2,683. Microarray MA-plots denoted concordance between expected and calculated mean and log-ratio intensity values for control spots (S1 Fig).

### Transcriptomic profile of transfected *A. aegypti*

A generalized transcriptomic profile of transfected female *Aedes aegypti* (Fig 1) was developed based on genes that were consistently upregulated in at least seven out of the eight DEG sets (Aa1-Aa8) derived from this organism (complete DEG sets in S1 Table).

**Fig 1.**
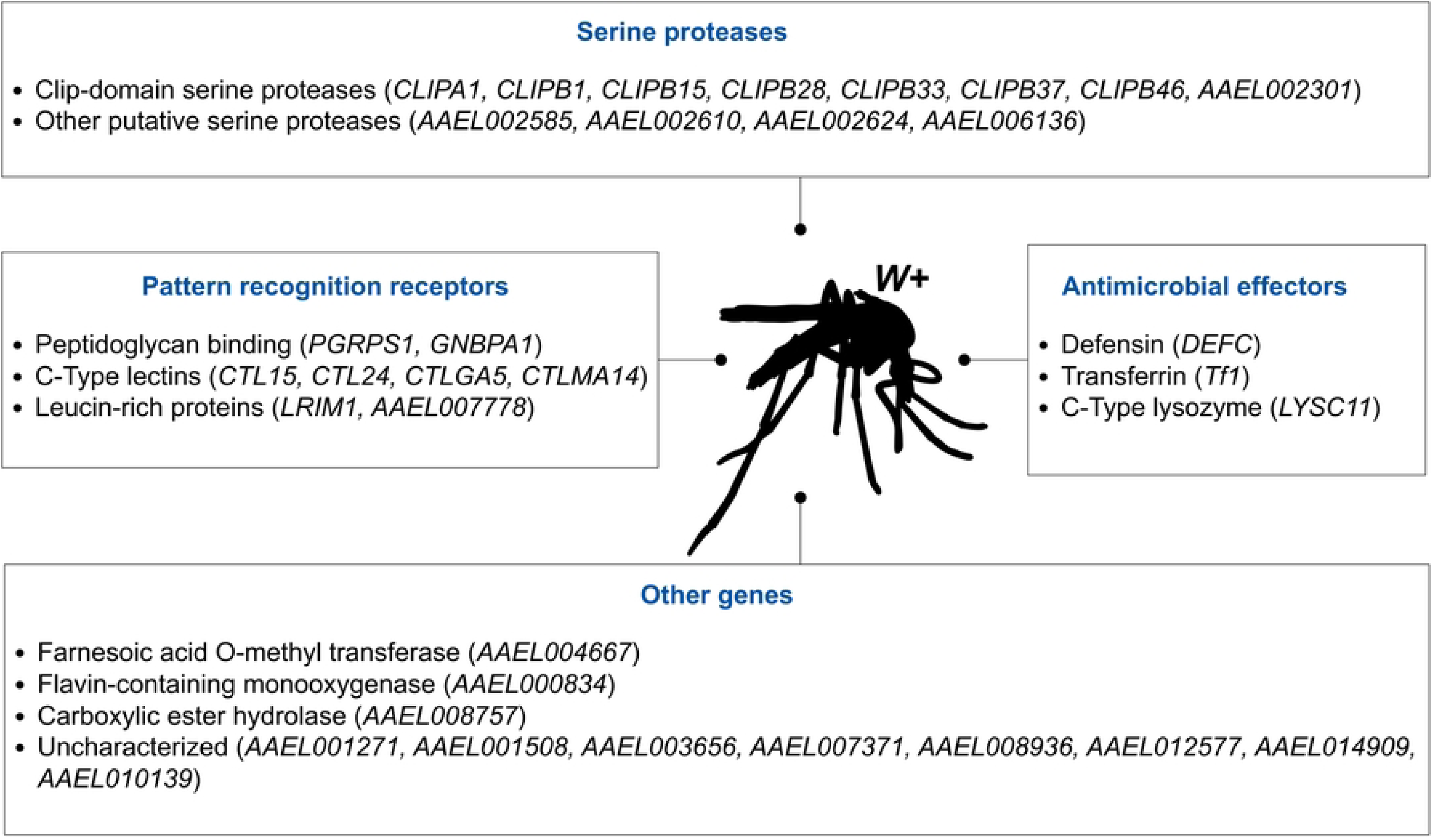
Genes that were consistently upregulated by *Wolbachia* transfections in female *A. aegypti* mosquitoes.

As shown in Fig 1, there were 34 consistently upregulated genes (referred to as CURGs in what follows). Clip-domain serine proteases, pattern recognition receptors and antimicrobial effectors comprise 19 of these genes, which are linked to the innate immune mosquito response. Immune priming is considered as one of the main mechanisms of viral blocking by *Wolbachia*, and concordantly, has received vast attention in the literature [34,35,63]. Our analysis contributes to this field of knowledge, by identifying a core of innate immune genes that are invariably upregulated by *Wolbachia* in *A. aegypti* females. More importantly, it also shows the consistent upregulation of genes that have been overlooked, identifying relevant study targets to deepen the mechanistic understanding of the *A. aegypti*-*Wolbachia* endosymbiotic system.

### Transcriptomic signature of *Aedes aegypti*’s immune priming by *Wolbachia*

Peptidoglycan binding proteins (*PGRPS1*, *GNBPA1*), defensin (*DEFC*), C-type lysozyme (*LYSC11*) and transferrin (*Tf1*) were CURGs in transfected female *A. aegypti* (Fig 1). All these genes encode hemolymphatic proteins related to the Toll pathway, which mainly responds to Gram-positive bacteria by activating the transcription of antimicrobial effector molecules [64]. Concretely, upon binding of peptidoglycan, the pattern recognition receptors *PGRPS1* and *GNBPA1* elicit a protease cascade, activating the Toll signaling pathway and thereby the transcription of effectors like the defensin *DEFC* or the lysozyme *LYSC11* [64,65]. Transferrins such as *Tf1* also have an innate immune function regulated by the Toll pathway, which can be induced to sequester iron from the hemolymph, limiting its availability for pathogens [66].

Our analysis is consistent with the well accepted fact that *Wolbachia* primes the Toll pathway in *A. aegypti* [34,35,63,67,68]. However, results show that only elements from the upstream and downstream of the Toll pathway appear to be consistently upregulated at the transcriptomic level, while many of the genes that perform the intracellular signal transduction are absent from all DEG sets (S1 Table). This is the case of the signaling elements *cact* (*AAEL000709*), *MYD* (*AAEL007768*), *Tube* (*AAEL007642*), *Pelle* (*AAEL006571*) and the transcription factor *rel1B* (*AAEL006930*), all of which were screened for in *A. aegypti*’s microarrays. Our analysis also suggests that downstream effectors *DEFC*, *LYSC11* and *Tf1* are preferentially induced by *Wolbachia*, while other canonical effectors of the mosquito Toll pathway such as cecropins and gambicin are present in no more than two DEG sets (S1 Table).

As *DEFC* is also inducible by the mosquito immunodeficiency (IMD) pathway, a broadly-conserved NF-κB immune signaling pathway that regulates a potent antibacterial defense response [69], the possibility of its upregulation being mediated by this pathway instead of Toll could be considered. However, we did not find any of the IMD pathway-activating receptors *PGRP-LA*, *PGRP-LE* or *PGRP-LD* to be upregulated in any of the DEG sets, while the inhibitor receptor *PGRP-LB* was upregulated in Aa2 and Aa5 (S1 Table). The upregulation of the IMD-pathway related transcription factor *REL2* in three DEG sets (Aa2, Aa5 and Aa8; S1 Table) offers the only transcriptomic evidence in favor of an activated IMD pathway, however, it does not explain the consistent upregulation of *DEFC* in most DEG sets, thus reinforcing the idea that the activation of Toll pathway is the one that is persistent in transfected *A. aegypti*.

Eight genes encoding putative Clip-domain serine proteases (CLIPs) were CURGs in transfected *A. aegypti* (Fig 1). Insect CLIPs are extracellular proteins that participate in protease signaling cascades upon injury or infection [70,71]. The consistent upregulation of eight CLIPs points towards a generalized activation of extracellular protease cascades, which in turn can be linked with two downstream immune responses: melanization mediated by prophenoloxidase and Toll pathways. Indeed, although the mapping between specific CLIPs, protease cascades, and targeted downstream processes is still in progress for *A. aegypti* [72–74], the fact that these two pathways are activated by protease cascades is well established. Furthermore, the CURG *CLIPB15* has been shown to participate in the upstream of the melanization and Toll pathways [73], while *AAEL002301* encodes a prophenoloxidase-activating factor 2-like Clip-domain (InterProScan: IPR041515), suggesting that it participates in the melanization process. These results are consistent with the evidence presented above showing that *Wolbachia* induction of the Toll pathway relies transcriptomically on upstream elements. They are also consistent with the priming of the melanization pathway that has been previously reported in *Wolbachia* infected hosts [75,76], and which is supported here by the joint overexpression of three prophenoloxidases *PPO4*, *PPO8* and *PPO10* in four DEG sets (Aa2, Aa4, Aa5 and Aa8; S1 Table).

Another four putative serine-type endopeptidases without CLIP-domain were CURGs in *A. aegypti*: *AAEL002585*, *AAEL002610*, *AAEL002624* and *AAEL006136* (Fig 1). Particularly, *AAEL006136*, may have immune functions, as it is responsive to ZIKV infection in *A. aegypti* [77] and it is a predicted ortholog of *Sp212*, which is upregulated after immune challenges and injury in *D. melanogaster* [78]. The remaining three genes encode putative trypsin domains (InterProScan: IPR001254), which is indicative of them being related to immune or digestive functions [79].

Four genes codifying C-Type lectins (CTLs) were CURGs in transfected *A. aegypti* (Fig 1). CTLs are proteins with a carbohydrate binding activity that generally depend on calcium ions, which are implicated in several aspects of insect innate immunity. CTLs act as pattern recognition receptors, being involved in agglutination, opsonization, encapsulation and melanization of microorganisms, as well as in the protection of the gut microbiome from the action of host antimicrobial peptides [80,81]. CTLs have also been implicated in facilitating infection of viruses and *Plasmodium* parasites in mosquitoes [81,82]. Concretely, *CTL15*, here identified as a CURG, has both a role in *A. aegypti*’s gut microbiome homeostasis, as well as in facilitating DENV infection in the same organism [80,82]. Also, *CTL24* was found to promote the infection of *A. aegypti* with the Japanese Encephalitis Virus [83]. Finally, *CTLMA14* has been identified as relevant for *A. aegypti*’s resistance to *E. coli*, while promoting DENV replication in the same host [80,82].

The two CURGs *LRIM1* and *AAEL007778* (Fig 1) are predicted to be leucine rich–repeat immune genes, a gene family that serve as receptors for mosquito innate immunity [84,85]. Particularly, *LRIM1* encodes a secreted protein that is antagonistic to *Plasmodium berghei* [86,87] and the bacteria *E. coli* in the mosquito *Anopheles gambiae* [88], as well as highly responsive to the arboviruses ZIKV and CHIKV in *A. aegypti* [85].

In summary, we identified a core of immune genes that are persistently upregulated by *Wolbachia* transfection in *A. aegypti*. Our results are consistent with *Wolbachia*’s priming of the Toll pathway in *A. aegypti* and show that, at the transcriptomic level, this phenomena is driven by the upregulation of genes in the upstream of the pathway: the pattern recognition receptors *PGRPS1* and *GNBPA1*, the CLIP-protease *CLIPB15* and possibly additional CLIP-proteases involved in extracellular signaling cascades. Our analysis also shows that the downstream effector genes that are preferentially upregulated by *Wolbachia* transfections are *DEFC*, *LYSC11* and *Tf1*, which are likely induced by the Toll-pathway and not the IMD pathway. C-type lectins and leucine rich-repeat immune genes compose the additional CURGs. Given the described roles of these genes in *A. aegypti*’s viral infection and their consistent upregulation by transfection here identified, we propose them as persistent key elements in the interface of host-*Wolbachia*-virus, with high relevance for unveiling the specifics of this interaction.

### Novel genes that are consistently upregulated by *Wolbachia*

*AAEL004667*, a gene which codifies for a farnesoic acid O-methyltransferase (FAMeT) domain, was consistently upregulated by *Wolbachia* in *A. aegypti* (Fig 1). FAMeT catalyzes the conversion of farnesoic acid to methyl farnesoate, which is a late step in the biosynthesis of insect juvenile hormones [89,90]. In *A. aegypti*, juvenile hormone III (JHIII) is a key regulator of development and reproduction, and its titer determines the expression of thousands of genes [91,92]. Although FAMeT may not participate in the preferred branch of JHIII biosynthesis in *A. aegypti* [93], this does not preclude its upregulation from influencing JHIII dynamics. FAMeT upregulation could affect JHIII biosynthesis by cooperating with the enzyme from the preferred branch (juvenile hormone acid methyltransferase or JHAMT), or by deviating the common substrate of these two enzymes to the alternative production of methyl farnesoate.

Liu et al. (2014) [94] proposed that paternal defects of *Wolbachia*-infected testes associated with cytoplasmic incompatibility were due to an alteration of JHIII concentration mediated by JHAMT upregulation. This suggests that an hypothetical effect of FAMeT upregulation in JHIII levels could also lead to cytoplasmic incompatibility and thus be of direct importance for *Wolbachia*-based arboviral control strategies. Additionally, It has been reported that methyl farnesoate can partially substitute the functions of JHIII in *A. aegypti* [95] and possibly serves additional uncharacterized functions [93], so that an upregulation of FAMeT could have deep consequences for mosquito physiology on this basis. *Wolbachia* interactions with the juvenile hormone biosynthetic pathway have been previously reported [94]. Particularly, a proteomic assay on transfected *A. aegypti* ovaries showed downregulation of farnesol dehydrogenase and mevalonate kinase, two enzymes in the JHIII biosynthetic pathway [96]. However, to the best of our knowledge, our study is the first showing *Wolbachia*’s consistent upregulation of the putative FAMeT codifying *AAEL004667* in *A. aegypti*. Another consistently upregulated gene was *AAEL000834*, which codifies a flavin-containing monooxygenase (FMO) (Fig 1). Monooxygenases are oxidoreductases that catalyze the incorporation of an oxygen atom from molecular oxygen to a compound, while FMOs are monooxygenases that specifically use flavin adenine dinucleotide (FAD) as a prosthetic group [97,98]. In mammals, FMOs serve to neutralize substances that are foreign to the normal functioning of the organism (i.e., *xenobiotics*) as well as to catalyze a step in the biosynthesis of taurine [99,100]. Insect FMOs are less known, although it is believed that they play an analogous role to mammal FMOs, and they have been shown to provide resistance to certain insecticides [97,101,102].

Our work identifies FMO (*AAEL000834*) as a relevant target of study for unveiling the fundamental mechanisms of *A. aegypti*-*Wolbachia* interaction, pointing out for the first time the persistence of its upregulation in this endosymbiotic system. Furthermore, the importance that understanding the *Wolbachia-AAEL000834* interaction has for strategies of arboviral disease control is emphasized by the association of this gene with arboviral susceptibility [101]. Indeed, Kumar et al. (2023) [101] suggest that the upregulation of *AAEL000834* may have an agonistic effect on CHIKV infection. This is not inconsistent with the anti-CHIKV blocking provided by *Wolbachia* [24,103], as this agonistic effect could be overcomed by several other CHIKV-antagonistic effects of *Wolbachia* in its host, such as immune priming and resource competition.

A putative carboxylesterase (*AAEL008757,* EC 3.1.1.1), was also consistently upregulated by *Wolbachia* (Fig 1). As carboxylesterase is a broad category, it is difficult to associate this transcriptomic effect with a particular process. Transcriptomic studies in *A. aegypti* have shown that the expression of *AAEL008757* varies according to several factors, such as age and sex [104], insecticide resistance [105], ZIKV infection [77] and time after mating [106], implying that it may have multiple physiological roles. Seven additional genes (*AAEL001271*, *AAEL001508*, *AAEL003656*, *AAEL007371*, *AAEL008936*, *AAEL012577* and *AAEL014909*) were consistently upregulated by *Wolbachia* (Fig 1), all of which are essentially uncharacterized. The last two of them are predicted to encode a DUF4789 domain, the function of which is unknown [79]. We propose that *AAEL008757* and these additional seven genes should be further characterized, as our analysis suggests that their upregulation is not a matter of chance but rather they could underlie characteristic endosymbiotic phenomena. It will be of particular interest to determine the functions performed by the DUF4789 domain, which is codified by two of these consistently upregulated genes.

### Patterns of *Wolbachia* gene-level effects across *Diptera* hosts

Next, we broaden our search by including DEG sets from native infections (Af, Dm1-Dm5, and Dp1-Dp4, from Table 1). In order to allow a direct comparison across species, all DEG sets were mapped to *D. melanogaster* orthologs, by which 3,302 unique genes were found contained in or mapped to DEG sets (S2 Table). Most of these genes (2,250) were exclusive to one DEG set, denoting an overall specificity of *Wolbachia* effects at the gene level. However, we identified ten genes that showed defined expression patterns across datasets (Table 2). Pattern A: genes consistently overexpressed on *A. aegypti* transfections, but underexpressed or unchanged on native infections; pattern B: genes frequently over/underexpressed on native infections but never on transfections; and pattern C: genes that were overexpressed across both types of infections and in several host taxa. To facilitate following discussions, we will refer to 1:1 *Drosophila*/*Aedes* ortholog pairs presented in Table 2, through the name of the *A. aegypti* gene, unless otherwise stated.

**Table 2.**
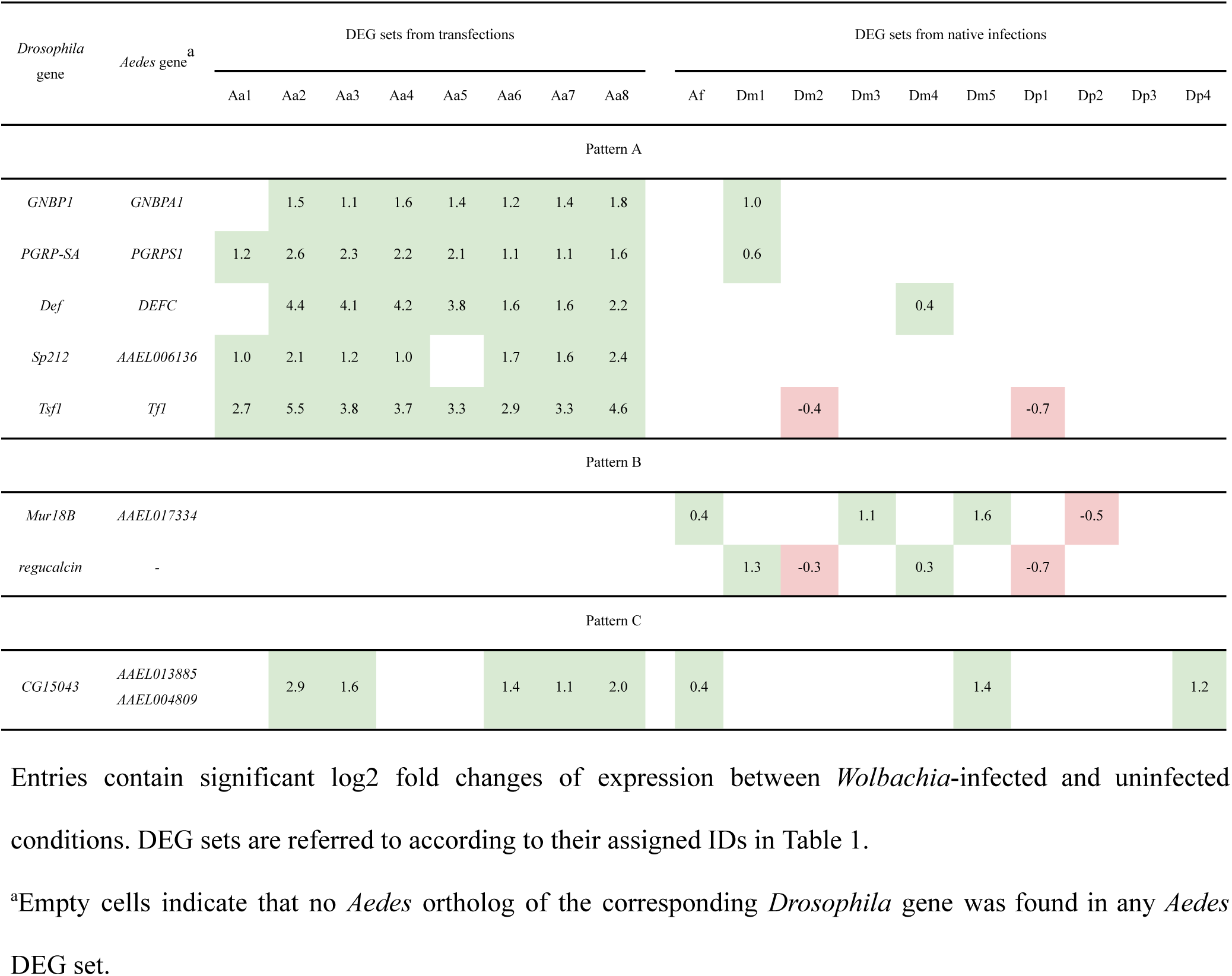
Expression patterns of DEGs.

### Five immune response genes lose responsiveness to *Wolbachia* over time (Pattern A)

Pattern A was displayed by five genes previously found to be CURGs in *A. aegypti* transfections, but that in native infections showed no differential expression (*AAEL006136*), infrequent differential expression (*GNBPA1*, *PGRPS1*, *DEFC*) or even downregulation (*Tf1*) (Table 2). Previous qRT-PCR assays have validated both the upregulation of *DEFC*, *AAEL006136* and *Tf1* in *A. aegypti* transfections [34,107] as well as the weakness of *GNBPA1*, *PGRPS1* and *DEFC* responses and the downregulation of *Tf1* in *D. melanogaster* native infections [34].

Four of the five genes displaying pattern A —*GNBPA1*, *PGRPS1*, *DEFC* and *Tf1*— are implicated in *A. aegypti*’s antiviral responses. As shown in the previous section, the upregulation of the peptidoglycan receptors *GNBPA1* and *PGRPS1* appears to be critical for *Wolbachia*’s activation of the host’s Toll pathway, which in turn has been shown to provide an anti-DENV mechanism to *A. aegypti* [65,108]. Additionally, silencing of *DEFC* has been shown to enhance DENV infection in *A. aegypti-wAlbB* [68] and the effect of *w*Mel in *DEFC* expression is strongly modulated by ZIKV/DENV coinfection in *A. aegypti* [109]. Transferrins provide insect tissues with iron [110], an element that mediates *Wolbachia*’s induction of oxidative stress, which is a key mechanism for arboviral blocking [68]. Particularly, Zhu et al. (2019) [111] showed that silencing of transferrins in *A. aegypti* enhanced DENV infection, while oral supplementation with the product of *Tf1* had the opposite effect. Additional evidence places transferrins at the interface of host, virus and *Wolbachia*, as *Tf1* was dynamically expressed following ZIKV inoculation [77] and *Wolbachia*’s effect in *Tf1* expression was strongly modulated by ZIKV/DENV coinfection [109].

Native host-*Wolbachia* systems have already undergone sustained coadaptation and thus can be taken as models for the future state of current transfections [13,20]. Under this paradigm, we propose that the expression pattern displayed by *GNBPA1*, *PGRPS1*, *DEFC*, *AAEL006136* and *Tf1* (Table 2) predicts the loss of their upregulation in the *A. aegypti*-*Wolbachia* systems currently established for arboviral control. Furthermore, the above mentioned contributions to antiviral defenses make it reasonable to expect that expression changes of *GNBPA1*, *PGRPS1*, *DEFC* and *Tf1* would undermine the viral blocking phenotype conferred by *Wolbachia*, and therefore lead to an increased risk of human disease in intervened areas. Acknowledging the risk of a weakening of these immune genes’ responses, one crucial question to address is if their lower responsiveness to native infections follows solely from reduced bacterial titer and tropism or reflect other qualitative differences. Answering this will help to determine appropriate ways to preserve these antiviral mechanisms over time. Taking all these into account, we recommend that *Wolbachia*’s upregulation of *GNBPA1*, *PGRPS1*, *DEFC* and *Tf1* in *A. aegypti* should be kept under study, given the direct contribution of these genes to the arboviral blocking phenotype.

### *Mur18B and regucalcin* are only responsive to native infections (Pattern B)

Differential expression of *Mur18B* and *regucalcin* orthologs was common among native *Wolbachia* infections, but these genes showed no significant effects in *A. aegypti* transfections (Table 2). This lack of regulation stands even when considering, additionally to g:Profiler ortholog assignment, all *A. aegypti*’s orthologs of *Mur18B* and *regucalcin* predicted by OrthoDB [61], which were effectively screened for in the corresponding microarrays. Although we did not find differential expression of *regucalcin* by native *Wolbachia* in *A. fluviatilis*, the analysis of the same dataset by Caragata et al. (2017) showed two differentially expressed transcripts that mapped to *regucalcin* orthologs (AAEL001020 and AAEL000757), a discordance we attribute to methodological differences (aggregation of RNA-Seq counts at the gene level rather than the transcript level). Taken together, these results indicate that *Mur18B* and *regucalcin* are regulated by *Wolbachia* in the three natively infected species studied, but not in transfected *A. aegypti*.

Given the observed pattern, one can speculate that the regulation of *Mur18B* and *regucalcin* reflect adaptive host-*Wolbachia* interactions. *regucalcin,* AAEL001020 and AAEL000757 are annotated to the Senescence marker protein-30 family (InterProScan: IPR005511), which activate or inhibit many calcium-dependent enzymes and regulate the intracellular calcium homeostasis, by controlling the activity of calcium channels and pumps [79,112,113]. In turn, *Mur18B* and AAEL017334 are predicted to bind chitin, a carbohydrate that serves critical structural roles in insects [58,114]. We will further link these observations to two ubiquitous functional effects of *Wolbachia*, i.e. calcium ion binding and chitinase activity.

### *CG15043* is upregulated in native infections and transfections, comprising all host taxa (Pattern C)

As shown in Table 2, *Drosophila*’s *CG15043* or its *Aedes’* orthologs *AAEL013885* and *AAEL004809* were upregulated both in native infections and transfections, comprising hosts from all four studied *Diptera* species. *CG15043*, *AAEL013885* and *AAEL004809* are mainly uncharacterized, however, they are predicted to encode domains of the MBF2 family of transcription activators, PFAM: PF15868 [115]. This is remarkable considering that *Wolbachia* exerts genome-wide transcriptional effects, in fact, our work found 3,302 *Drosophila* genes mapped to DEG sets, which is around one quarter of all *D. melanogaster* protein coding genes [116]. The overexpression of putative MBF2 transcription activators across several different host-*Wolbachia* combinations puts these genes as novel targets for understanding the mechanisms underlying *Wolbachia*’s wide gene regulation capabilities.

### Patterns of *Wolbachia* functional effects across *Diptera* hosts

We searched for transcriptomic patterns that may not appear at the level of individual genes but rather of sets of genes with common functions or localizations. After grouping DEG sets according to direction of regulation, enrichment analyses of GO terms were performed on each subset with an exploratory FDR *q*-value cutoff of 0.20. 4,948 GO terms were found to be enriched (S3 Table), most of which (2,738) were present in just one subset, showing again an overall specificity of *Wolbachia*’s effects across biological conditions. Table 3 presents the enrichment pattern of the five most frequently enriched GO terms, as well as of the additional terms calcium ion binding and chitinase activity, on which we focused given their relation to *regucalcin* and *Mur18B* (previously found as commonly regulated in native infections) and which were also commonly regulated across DEG sets. In what follows, we refer to a functional term as up/downregulated in a DEG set if it was enriched in the subset of up/downregulated genes of that DEG set, as presented in Table 3.

**Table 3.** Enriched GO terms across DEG sets.

| GO Term |  | DEG sets from transfections |  |  |  |  |  |  |  | DEG sets from native infections |  |  |  |  |  |  |  |  |  |
| --- | --- | --- | --- | --- | --- | --- | --- | --- | --- | --- | --- | --- | --- | --- | --- | --- | --- | --- | --- |
|  |  | Aa1 | Aa2 | Aa3 | Aa4 | Aa5 | Aa6 | Aa7 | Aa8 | Af | Dm1 | Dm2 | Dm3 | Dm4 | Dm5 | Dp1 | Dp2 | Dp3 | Dp4 |
| Extracellular region | + | 10 | 24 | 7 | 12 | 17 | 30 | 26 | 55 | 4 | 31 | 37 | 23 | 60 | 4 | 2 | 10 | 5 | 40 |
|  | - |  | 1 |  | 2 |  |  |  | 3 | 2 | 31 | 24 | 13 | 7 | 2 | 56 | 70 | 22 | 4 |
| Serine-type endopeptidase activity | + | 17 | 38 | 18 | 27 | 33 | 80 | 74 | 114 | 6 | 6 | 23 | 10 | 14 |  |  | 2 | 1 | 4 |
|  | - | 1 | 1 | 1 | 3 |  | 1 |  | 3 | 5 | 5 | 4 | 4 |  |  | 7 | 15 | 5 | 2 |
| Heme binding | + | 2 | 19 |  | 2 | 5 | 11 | 16 | 37 | 4 | 9 | 4 | 6 | 7 | 1 |  | 2 |  | 3 |
|  | - |  | 1 |  | 1 | 3 |  | 1 | 7 | 1 |  | 5 | 3 | 2 | 1 | 11 | 7 | 11 |  |
| Mono-oxygenase activity | + | 3 | 18 |  | 2 | 4 | 9 | 13 | 35 | 3 | 10 | 5 | 5 | 7 | 1 |  | 2 |  |  |
|  | - |  |  |  | 1 | 4 |  |  | 4 |  |  | 2 | 4 | 1 | 1 | 9 | 11 | 8 |  |
| Endopeptidase inhibitor activity | + | 2 | 7 | 1 | 4 | 3 | 14 | 11 | 22 |  | 6 | 6 |  | 5 |  |  |  |  | 4 |
|  | - |  |  |  |  |  |  |  |  | 1 |  | 4 | 2 |  |  | 4 | 4 | 2 |  |
| Calcium ion binding | + | 1 | 16 | 1 | 4 | 5 | 8 | 6 | 19 | 1 | 2 | 11 | 2 | 11 | 2 |  | 10 | 2 | 16 |
|  | - | 1 |  |  | 3 |  |  | 1 | 6 | 3 | 5 | 3 | 2 | 2 | 3 | 3 | 2 | 3 | 2 |
| Chitinase activity | + |  |  |  |  | 1 | 1 | 1 | 1 |  | 2 |  | 1 |  |  |  | 1 |  | 1 |
|  | - |  |  |  |  |  |  |  |  |  |  | 2 |  | 1 |  | 1 | 2 | 1 |  |

Entries show the number of upregulated (+) or downregulated (-) genes in a given DEG set that were annotated with a given functional term. Green or red cells indicate significant enrichment of a functional term in a given subset of up and downregulated genes (respectively), with FDR q-value cutoff of 0.20. The first five rows correspond to the five most frequently enriched GO terms.

### Extracellular region, serine-type endopeptidase and endopeptidase inhibitor activities

Extracellular region (GO:0005576), serine-type endopeptidase activity (GO:0004252) and endopeptidase inhibitor activity (GO:0004866) were among the five most frequently enriched GO terms (Table 3). Concerning *A. aegypti*, the common upregulation of extracellular region and serine-type endopeptidase activity is consistent with our transcriptomic profile based on individual CURGs, which included multiple (CLIP and non-CLIP) serine-type endopeptidases and other extracellular components of the innate immune system. Additionally, the common upregulation of endopeptidase inhibitor activity revealed by enrichment analyses complements our generalized transcriptomic portrait of transfected *A. aegypti* and should reflect the activity of serine-protease inhibitors that constrain the spatial and temporal extension of extracellular protease cascades [70,71].

The regulation of the three previously mentioned GO terms is also common among native infections, although the direction of the regulation is not strictly positive as in *A. aegypti* transfections. Natively infected *A. fluviatilis* show upregulation of extracellular region genes, both up and downregulation of serine-type endopeptidase activity and it does not show regulation of endopeptidase inhibitor activity (set Af; Table 3). The significance and direction of regulation of these GO terms is variable across DEG sets from *D. melanogaster*-*w*Mel (sets Dm1 to Dm5; Table 3), although upregulated genes associated with these terms usually exceed in number to the downregulated ones, denoting an overall tendency towards upregulation. Finally, *D. paulistorum*-*w*Pau males show downregulation of the three GO terms (sets Dp1 and Dp2; Table 3), while females show no clear preferential direction (sets Dp3 and Dp4; Table 3).

Activity of serine-type endopeptidases and endopeptidase inhibitors is linked to viral blocking in transfected *A. aegypti*, as they mediate extracellular protease cascades which trigger the Toll pathway and the expression of antiviral effectors, as previously exposed. Moreover, we showed that *Wolbachia*’s priming of the Toll pathway in *A. aegypti* strongly relies on the upregulation of these upstream events. Therefore, it is necessary to consider the loss of these effects upon coadaptation of *A. aegypti*-*Wolbachia* systems. The clearest contrast with *A. aegypti* trends regarding the regulation of these genes comes from *D. paulistorum*, where downregulation predominates. However, *D. paulistorum* native infection may be not the best model for coadapted *A. aegypti*-*Wolbachia*, as the first one constitutes an extreme case of obligate mutualism where hosts cannot reproduce without *Wolbachia* [117]. In such obligate relationships, strong selective pressures favor tolerance over resistance, that is, hosts are strictly selected to limit the negative consequences of *Wolbachia* infection without limiting the infection itself [118].

Facultative symbioses such as those established by *A. fluviatilis* and *D. melanogaster* should better reflect the result of coadaptation when selection of tolerance traits is not imperative. As previously discussed, DEG sets from these systems show an overall tendency towards upregulation of serine-type endopeptidases and endopeptidase inhibitors, although to a lesser degree than DEG sets from *A. aegypti*, occasionally with up and downregulation occurring simultaneously. These observations suggest that a partial loss of the upregulation of these groups of genes is possible. Further evidence on this possibility is provided by DEG sets Aa6, Aa7 and Aa8, which were derived from three *A. aegypti*-*w*Mel populations introgressed in Australia in 2011, 2013/14 and 2017, respectively [26]. As shown in Table 3, the number of serine-type endopeptidases and endopeptidase inhibitors were considerably higher in set Aa8 than in Aa6 and Aa7, indicating that these effects decline to some degree when transfected mosquitoes are exposed to wild conditions.

The importance of *A. aegypti*’s upregulation of serine-type endopeptidases and endopeptidase inhibitors for arboviral control and the prospect of its future reduction highlights the necessity of monitoring the effects of *Wolbachia* on these genes over time and to design strategies to prevent or react to these changes. Also, our previous discussions imply the importance of preventing the *A. aegypti*-*Wolbachia* system from acquiring an obligate character, as this may force the loss of resistance traits such as immune activation. In this regard, monitoring the appearance of dependence traits in *A. aegypti* (i.e., losses of host functions that become fulfilled by *Wolbachia*) would be relevant, as they are considered precursors for obligate mutualisms [118].

### Heme binding and monooxygenase activity

Two additional GO terms among the most frequently enriched were heme binding (GO:0020037) and monooxygenase activity (GO:0004497) (Table 3). These terms showed coupled enrichment patterns across DEG sets, which is due to the fact that both of them share differentially expressed genes, mostly cytochromes p450 (CYPs), a wide family of heme-dependent monooxygenases (S1 and S2 Tables). Caragata et al. (2017) [13] pointed out the commonality of the regulation of CYPs across *A. fluviatilis-w*Flu and *A. aegypti* transfections. Here, we extend the comparison to *D. melanogaster* and *D. paulistorum*, highlighting that CYPs are also commonly regulated in these systems. As shown in Table 3, the regulation of heme binding and monooxygenase activity in *A. aegypti*, *A. fluviatilis* and *D. melanogaster* is predominantly in the positive direction, while the opposite trend is observed in *D. paulistorum* (sets Dp1, Dp2 and Dp3; Table 3).

Regulation of CYPs can be linked to a broad-targeting mechanism of viral blocking by *Wolbachia*, which is the induction of reactive oxygen species (ROS). ROS induction by *Wolbachia* has been reported in several *Diptera* hosts, whereas evidence of this phenomena mediating *Wolbachia*’s viral blocking has been shown for transfected *A. aegypti* and natively infected *D. melanogaster* and *D. simulans* [68,119,120]. ROS induction by *Wolbachia* has been previously ascribed to electron leakage from the bacterial oxidative phosphorylation but also to the activity of host NADPH oxidases and dual oxidases, as some of these genes were strongly upregulated in *A. aegypti*-*w*AlbB [68]. However, these genes were almost absent from the analyzed DEG sets (S1 and S2 Tables), suggesting that, if there is some general underlying mechanism of ROS induction apart from *Wolbachia*’s central metabolism, it likely does not depend on these genes. On the other hand, our results show that a common transcriptomic trait of *Wolbachia*-infected *A. aegypti* and *D. melanogaster* is the upregulation of CYPs, the activity of which can constitute a major source of ROS in animal cells [121]. This allows to propose the upregulation of CYPs as a better explanation for the commonality of ROS induction by *Wolbachia* in *Diptera* hosts and, potentially, as a key trait for viral blocking.

### Calcium ion binding

As shown in Table 3, calcium ion binding (GO:0005509) genes were enriched in DEG sets from all four host species. Particularly, this molecular function was enriched in DEG sets from transfected *A. aegypti* (sets Aa2, Aa4) and natively infected *A. fluviatilis* (set Af), *D. melanogaster* (sets Dm2, Dm4 and Dm5) and *D. paulistorum* (sets Dp2, Dp4). Collectively, these results suggest that the regulation of calcium ion binding genes is a common phenomena in *Diptera*-*Wolbachia* associations. Although previous studies have found differential expression of calcium ion binding genes [13,26,28,29,31–35,62], this is the first work that highlights the ubiquity of these effects by *Wolbachia*.

It has been previously described that *Wolbachia* obtains its external vacuolar membrane by sequestering it from the Golgi apparatus and the endoplasmic reticulum, a process that implies a physical interaction and modifies the composition and organization of these organelles [122,123]. We speculate that the common transcriptomic alterations of calcium ion binding genes here shown could be reflecting this interaction, and that this causal link occurs via disruption of intracellular calcium ion gradients. Indeed, both the Golgi apparatus and the endoplasmic reticulum are intracellular calcium reservoirs, so it is reasonable to expect that their direct interaction with *Wolbachia* would modify intracellular calcium gradients with respect to uninfected cells. Furthermore, given the implications of calcium signaling in many critical cellular processes and the toxicity of calcium when exceeding its normal temporal and spatial boundaries [112,113,124], the shared transcriptomic regulation of calcium ion binding genes could be well explained as host compensatory responses to disruption of intracellular calcium gradients. In particular, the common native *Wolbachia*’s effects on *regucalcin* expression here described could constitute an adaptive host mechanism to restore calcium homeostasis, as *regucalcin* is a key regulator of the intracellular calcium distributions, which controls the activity of calcium channels and pumps, as well as of calcium-dependent enzymes [112,113].

Overall, given the centrality of calcium for biological phenomena, we propose that studying calcium related host-*Wolbachia* interactions is crucial for developing a solid mechanistic understanding of the endosymbiotic system.

### Chitinase activity

Chitin is a polysaccharide composed of N-acetylglucosamine (GlcNAc) residues with beta-1,4 O-glycosidic bonds, which is a fundamental component of the insect exoskeleton, internal linings such as the trachea and salivary ducts, the peritrophic matrix that covers the midgut epithelium, and the serosal cuticle that covers mosquito eggs [125–129]. As shown in Table 3, chitinase activity (GO:0004568) was significantly enriched in DEG sets from transfected *A. aegypti* (sets Aa5-Aa7), native *D. melanogaster*-*w*Mel (sets Dm1, Dm2), and native *D. paulistorum*-*w*Pau (set Dp2). Also, at least one chitinase was differentially expressed in two additional DEG sets from *D. melanogaster*-wMel (sets Dm3, Dm4) and in the three remaining DEG sets from *D. paulistorum* (sets Dp1, Dp3 and Dp4). Although these last effects were non-significant according to the enrichment analysis, it must be considered that chitinase activity comprises only 15 genes in *D. melanogaster* (compared *e.g.* to serine endopeptidase activity, that has 227 genes in the same organism) and, therefore, the relatively low number in Table 2 might be of biological relevance.

Furthermore, although in *A. aegypti* there is only one gene annotated with the GO term chitinase activity (AAEL013262), there are several more putative chitinases. Specifically, AAEL002023, AAEL003066 and AAEL001965 are all assigned to the InterProScan family IPR011583 (Chitinase II) and the Panther family PTHR11177 (Chitinase), while being annotated with chitin binding activity and combinations of additional GO terms such as hydrolysis of O-glycosyl compounds and chitin metabolic processes. Assuming that these genes indeed have chitinase activity, the alteration of this function by *Wolbachia* turns out to be even more common in *Aedes* hosts. In fact, we found an orthologous gene of AAEL003066 to be upregulated in *A. fluviatilis-w*Flu (set Af); as well as both AAEL002023 and AAEL013262 in female *A. aegypti*-*w*MelPop heads (set Aa5); AAEL002023 in female *A. aegypti*-wMelPop muscles (set Aa4); and both AAEL003066 and AAEL01965 in female *A. aegypti*-wMelPop whole bodies (set Aa2) (S1 Table). Additional literature supports the upregulation of chitinases by *Wolbachia* transfections. Geoghegan et al. (2017) [130] found upregulation of AAEL013262 and AAEL003066 at the protein level in *A. aegypti*-*w*Mel midguts, while Detcharoen et al. (2021) [33] found overexpression of a chitinase in *Drosophila nigrosparsa* when transfected with *w*Mel. Together, these observations indicate that the regulation of chitinases is an ubiquitous trait of host-*Wolbachia* systems, and that this regulation occurs strictly in a positive direction in the case of transfections.

We found only a few works that gave attention to the *Wolbachia*-chitin interaction. Quek et al. (2022) [131] showed that *Wolbachia* upregulates chitinase activity in the early stages of the filarial nematode *Brugia malayi*, and that this effect is necessary for the nematode to undergo exsheathment and escape the midgut barrier of its mosquito host. Sigle & McGraw (2019) [127] proposed chitin-related processes as a non-canonical point of interaction between *Wolbachia* and arboviruses. Caragata et al. (2017) [13], which found a putative chitinase to be upregulated in their *A. fluviatilis*-wFlu dataset, speculated that this regulation could be contributing to *Plasmodium* enhancement by disturbing the integrity of the peritrophic matrix. However, to our knowledge, no previous works have illustrated how generalized *Wolbachia* regulation of host chitinases is. Furthermore, we believe that this interaction should be explored further, for its implications in critical phenotypes for arboviral disease control, as depicted in Fig 2 and explained in the following.

**Fig 2.**
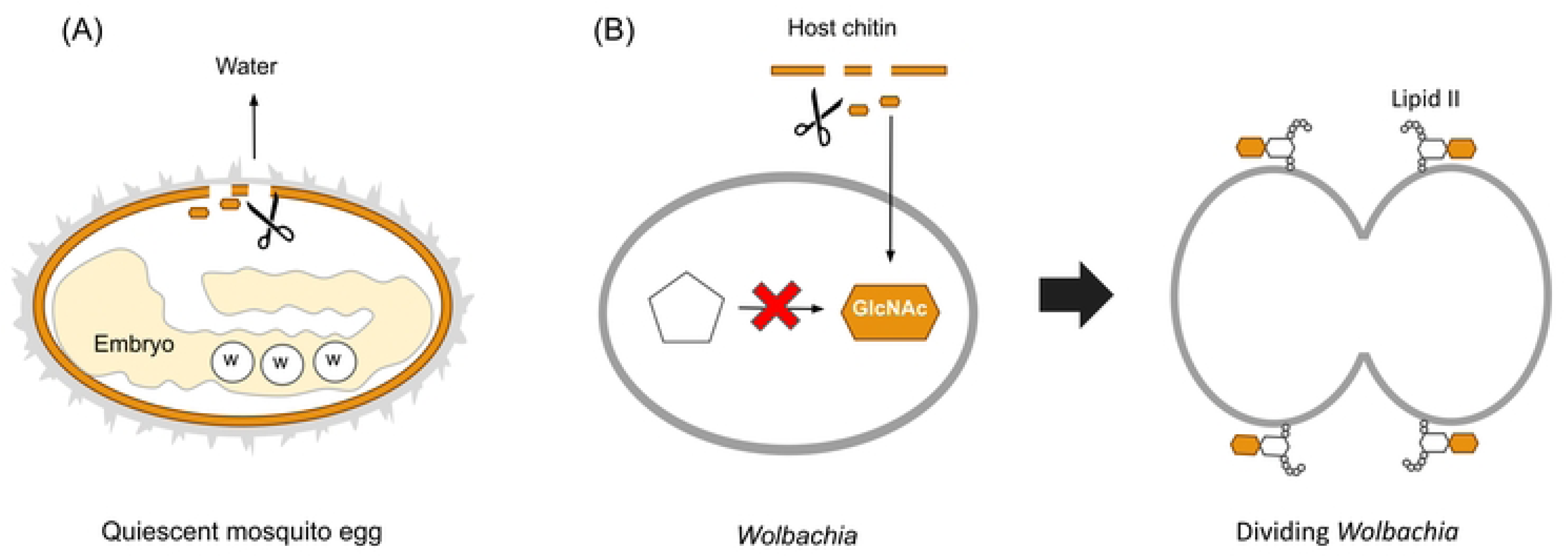
Associations proposed in this work between the observed ubiquity of chitinase regulation and critical phenotypes for vector control. (A) Upregulation of host chitinases by *Wolbachia* may be implicated in the loss of viability of quiescent eggs in transfected *A. aegypti*, by disrupting the chitinous serosal cuticle that provides them with desiccation resistance. (B) Chitinase regulation by *Wolbachia* may influence bacterial density in certain conditions by determining availability of GlcNAc, which is apparently unable to be synthesized *de novo* by *Wolbachia* despite needing it for lipid II synthesis (necessary for its cellular division). Scissor icons represent chitinases. Orange stripes and hexagons represent chitin and GlcNAc, respectively. Circles demarcated with a “W” denote *Wolbachia* cells within the mosquito embryo.

We claim there is a link between the consistent upregulation of chitinases and the loss of quiescent egg viability in transfected *A. aegypti*, which have been described as the main fitness cost of transfections and an obstacle for the scalability of *Wolbachia*-based arboviral control [132]. Indeed, mosquito quiescent egg survival strongly depends on desiccation resistance [132,133], which is both diminished by *Wolbachia* transfection [134] and strongly correlates with chitin content of the egg’s serosal cuticle [126,133]. Thus, it is reasonable to hypothesize that the chitinase upregulation unveiled in this work, triggered by transfected *Wolbachia,* weakens the serosal cuticle’s chitin, diminishing egg resistance to desiccation. *Wolbachia*-dependent exsheathment of *B. malayi* microfilariae, which is mediated by chitinase upregulation [131], provides an antecedent of such a *Wolbachia*-induced degradation of a host’s chitinous external structure.

We propose that chitinase regulation may directly influence *Wolbachia’s* titer, which is consistent with the fact that UDP-GlcNAc, an intermediary of synthesis of lipid II, is required for the coordination of its cellular division [135,136]. Despite its essentiality, it is not clear whether *Wolbachia* has the capability to synthesize GlcNAc *de novo* from fructose-6-P as some extracellular bacteria do [137,138], or if it must obtain it from the host, since it has not been reported in literature how this bacteria acquires GlcNAc [135,136,139]). Supporting the latter, *de novo* synthesis of GlcNAc in *Wolbachia* is absent in the KEGG database [137]. Furthermore, this metabolic dependence on host resources would not be an exception, as *Wolbachia* depends on several host amino acids and various metabolic gaps have been previously identified [140–143]. Therefore, we hypothesize that *Wolbachia* may induce chitinase activity to free and use host GlcNAc, as is the case for *Sodalis glossinidius*, the endosymbiont of the Tsetse fly [144].

If sequestration of host GlcNAc is the mechanism by which *Wolbachia* obtains this metabolite, it is reasonable to suppose that chitinase activity favors *Wolbachia*’s titer under certain conditions, such as nutrient abundance. This hypothesis is consistent with the expression patterns highlighted in this work (Table 3), particularly strict upregulation in transfections where *Wolbachia* is present in high titers, and variable direction of regulation in native infections, characterized by lower titers. As *Wolbachia* titer is positively correlated with critical traits such as viral blocking and fidelity of maternal transmission [22,23,25], we suggest that studying its dependence on a specific molecular activity such as chitinase would provide valuable gene targets for preserving the efficacy of *Wolbachia*-based vector control strategies.

As previously exposed, *Mur18B* or its *Aedes* ortholog *AAEL017334* were differentially expressed in the three natively infected hosts here studied, but not in transfections, suggesting that their responsiveness to *Wolbachia* reflects an adaptive mechanism. These are putative chitin binding genes and, particularly, *Drosophila*’s *Mur18B* is a predicted mucin, i.e., a type of gene that modulates the integrity of extracellular chitinous matrices in insects [145]. Thereby one can further speculate that common regulation of these genes in native infections reflects host adaptation to chitinase regulation by *Wolbachia*.

The commonality of *Wolbachia*’s chitinase regulation here observed, plus its putative role in *Plasmodium* enhancement [13], loss of quiescent egg viability and bacterial titer pose it as a main target for further research, as these phenotypes are critical for the safety, sustainability and scalability of *Wolbachia*-based arboviral control.

## Conclusions

Keeping the success of *Wolbachia*-based arboviral control requires a deep mechanistic knowledge of *Diptera*-*Wolbachia* systems, as well as anticipating the evolutionary path of transfections. By analyzing publicly available transcriptomic datasets, we were able to derive new hypotheses and targets of study that will help to broaden the understanding of *Diptera*-*Wolbachia* systems in general and *A. aegypti*-*Wolbachia* transfections in particular. This approach allowed to pinpoint gene-level and functional effects of *Wolbachia* that are consistent across transfected *A. aegypti* and are implicated in phenotypes critical for arboviral control, but that are likely to be lost after a coadaptation period, hence motivating their further study and monitoring over time.

## Supporting information

**S1 Fig. MA-plots of microarray datasets.** Scatter plots of M vs A for each microarray dataset, where M is the base-2 logarithm of the red/green intensity ratio of a spot, and A its average intensity. Control spots for M or A were identified according to platform descriptions and colored. Magenta: negative 3-fold change control. Blue: negative 10-fold change control. Orange: positive 3-fold change control. Red: positive 10-fold change control. Yellow: non-differentially expressed control. Purple: bright control. Gray: dark control.

**S1 Table. Complete DEG sets.**

**S2 Table. DEG sets mapped to *D. melanogaster* orthologs.** Log2 fold changes of DEGs mapped to *D. melanogaster* orthologs.

**S3 Table. Enrichment pattern of GO terms.** Number of DEGs annotated to each GO term in each upregulated (+) or downregulated (-) gene subset, and significance of the corresponding enrichment test. ‘Frequency’ denotes the number of upregulated or downregulated gene subsets that were enriched with a given GO term. ‘Term size’ denotes the size of a given GO term in *A. aegypti*.

***: padj < 0.05

**: padj < 0.10

*: padj < 0.20

