## Supplementary figures and images for "A look into the future: Using a transcriptomic meta-analysis of *Diptera*-*Wolbachia* systems to project the sustainability of arboviral control strategies"

### S1_Fig.tif

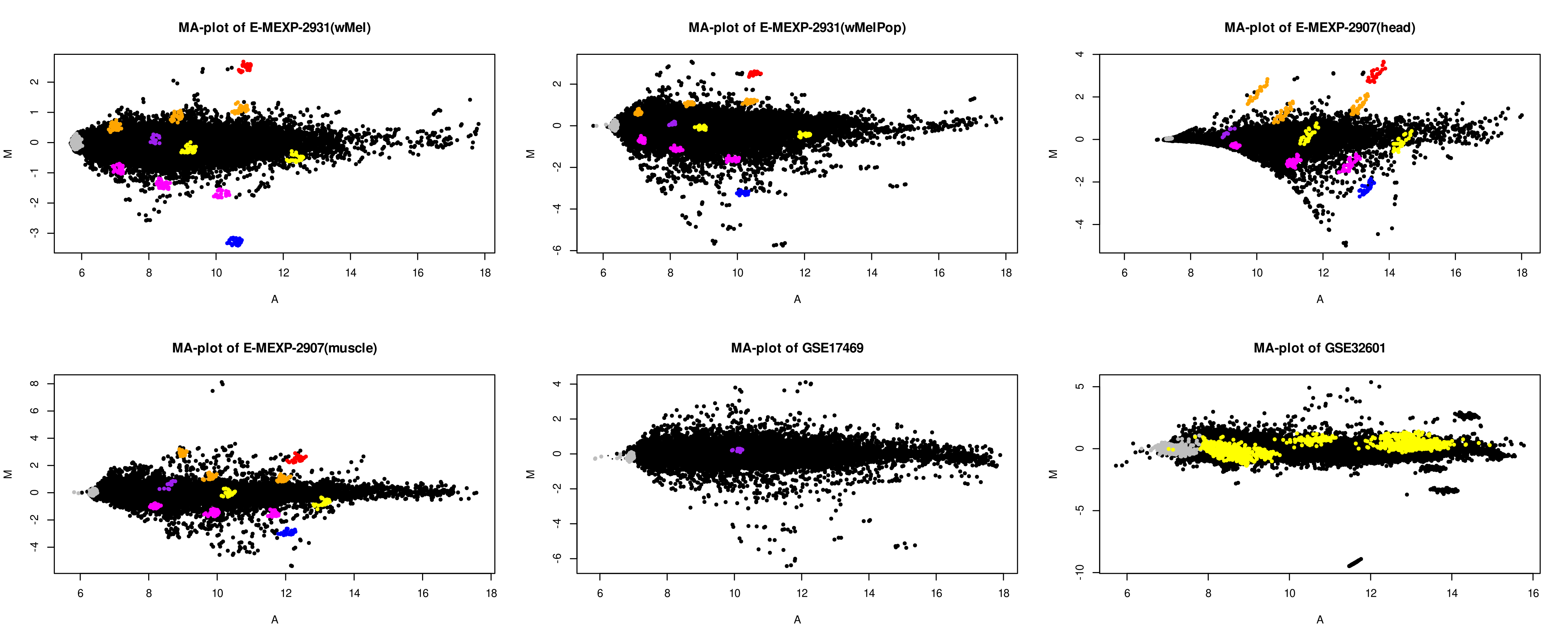
